# CardiacProfileR: An R package for extraction and visualisation of heart rate profiles from wearable fitness trackers

**DOI:** 10.1101/324004

**Authors:** Djordje Djordjevic, Beni K. Cawood, Sabrina K. Rispin, Anushi Shah, Leo H. H. Yim, Christopher S. Hayward, Joshua W. K. Ho

## Abstract

A person’s heart rate profile, which consists of resting heart rate, increase of heart rate upon exercise and recovery of heart rate after exercise, is traditionally measured by electrocardiography during a controlled exercise stress test. A heart rate profile is a useful clinical tool to identify individuals at risk of sudden death and other cardiovascular conditions. Nonetheless, conducting such exercise stress tests routinely is often inconvenient and logistically challenging for patients. The widespread availability of affordable wearable fitness trackers, such as Fitbit and Apple Watch, provides an exciting new means to collect longitudinal heart rate and physical activity data. We reason that by combining the heart rate and physical activity data from these devices, we can construct a person’s heart rate profile. Here we present an open source R package CardiacProfileR for extraction, analysis and visualisation of heart rate dynamics during physical activities from data generated from common wearable heart rate monitors. This package represents a step towards quantitative deep phenotyping in humans. CardiacProfileR is available via an MIT license at https://github.com/VCCRI/CardiacProfileR.

## Introduction

A heart rate profile is constructed from continuous measurement of heart rate before, during and after exercise, typically in the context of an exercise stress test. It contains important clinical parameters such as resting heart rate, maximal heart rate during exercise, and decrease of heart rate in the recovery phase. An increase in resting heart rate, a decrease in heart rate elevation in response to exercise, and a delay in heart rate recovery are significant predictors of sudden death (Cole et al., 1999; Jouven et al., 2005). These measurements, when derived from a controlled exercise stress test regime, have high intra-individual reproducibility over three years, indicating their usefulness as a diagnostic tool (Orini et al., 2017).

Heart rate parameters and profiles have been measured and constructed with exercise stress tests under controlled environments. This is often performed with a treadmill in a laboratory setting. With increasingly widespread availability of wearable wrist-worn fitness trackers, such as Fitbit and Apple Watch, we hypothesise that we can construct heart rate profiles for an individual using heart rate and physical activity (*e.g.*, as measured by global positioning system and accelerator) data from a person’s wearable monitor. The main advantage of this approach is that it opens unprecedented opportunity to continuously and non-invasively monitor a person’s heart rate profile under free-living conditions, which would better match this person’s realistic activity profile. These devices are socially acceptable, generally inexpensive, and are already widely available in many communities. This potential cardiac function monitoring technology can be seamlessly incorporated into a user’s daily life, without the need for clinically-based exercise stress testing.

A number of recent studies have assessed the accuracy of popular wrist-worn wearable fitness trackers (Dooley et al., 2017; Shcherbina et al., 2017). They found that while energy expenditure estimations are often less accurate, the measurements of heart rate are generally considered accurate. These findings suggest that it should be possible to extract heart rate dynamics information before, during and after an exercise event, which can be easily defined without accurate measurement of energy expenditure.

Here we present a new open source R package called CardiacProfileR. It is designed to efficiently extract heart rate and physical activity data from a Training Center XML (TCX) file which is generated by most modern fitness tracking devices. CardiacProfileR can identify periods of active exercise from the data, and construct a heart rate profile for each period of active exercise. This package produces interactive visualisation of one or multiple heart rate profiles in two dimensions or three dimensions. This enables aggregation and comparison of data from multiple periods of exercise or comparison between multiple people. As far as we know, CardiacProfileR is the first open source R package that provides an end-to-end analysis framework for wearable fitness sensor data.

## Program features

CardiacProfileR can be installed directly from the Github repository using the devtools R package. A comprehensive demo and a collection of example data files are included in the repository to illustrate the package’s functionalities. TCX files can be loaded individually or as multiple files in a folder. The data from each file are stored in a data frame after pre-processing to remove common issues with the data such as measurement gaps or time interval variations. Our package allows users to identify periods of active exercise (Fig. 1A). Activities are found by identifying when physical movement is not zero for an extended period of time, with a user-adjustable parameter allowing for short breaks (e.g. waiting at a traffic light).

**Figure 1.**
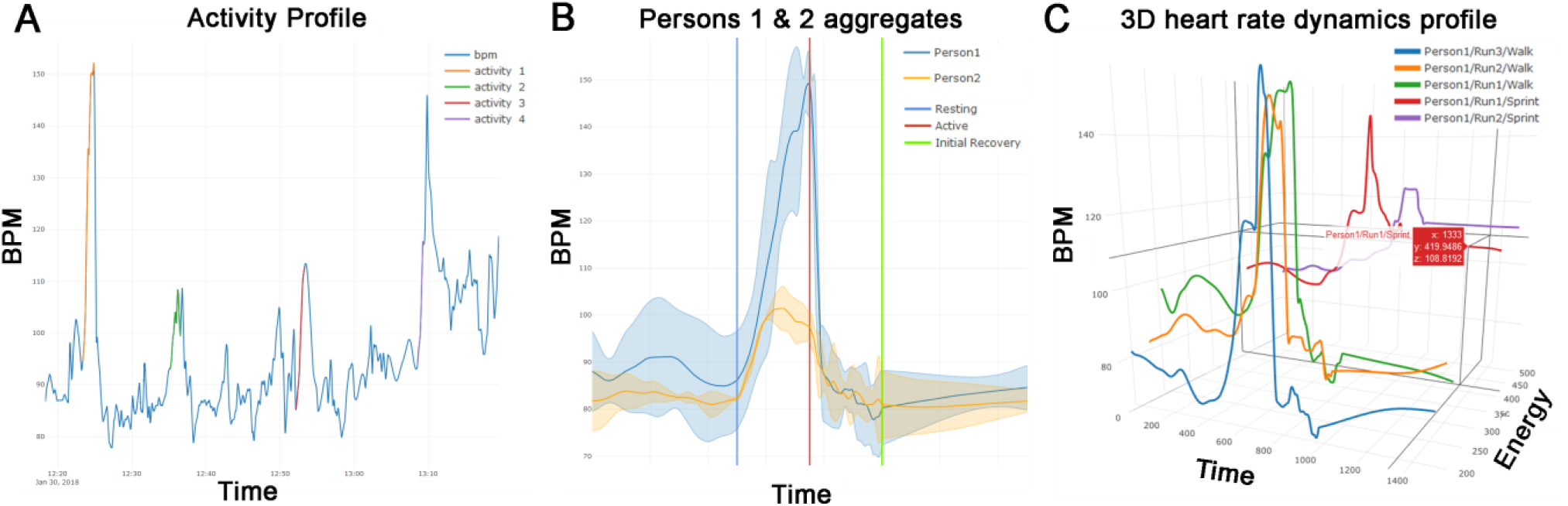
Extraction and visualisation of heart rate dynamics from heart rate and physical activity data from a Fitbit fitness tracker. **(A)** CardiacProfileR uses physical activity data to determine periods of active exercise. In this plot, four periods of exercise were identified and were highlighted on the heart rate time course. **(B)** Multiple physical exertion events of the same person can be summarised by an average scaled heart rate profile, showing the relationship between heart rate (in beat per minute; BPM) and time before, during (scaled) and after a period of exercise. This two-dimensional (2D) plot shows the average heart rate profile of two individuals upon performing the same exercise. **(C)** A three-dimensional (3D) plot showing multiple heart rate profiles from the same person from multiple periods of exercises based on their estimated energy expenditure.

Once the activities are found, a heart rate profile is created for each activity. A profile consists of heart rate measurements for four scaled temporal periods: the pre-activity resting period, the activity period, the initial recovery period and the final recovery period (Fig. 1B). An aggregate profile can be created for each person to visualise the variability between heart rate profiles generated from multiple similar activities. The aggregate profile is useful for comparing the shape and variability of heart rate profiles of the same person or across multiple people (Fig. 1B).

To compare multiple activities of one person with different energy expenditure (e.g., walking vs. running vs. sprinting), multiple heart rate profiles can be visualised in a three-dimensional plot (Fig. 1C) or surface plot, with the third axis being the inferred maximum energy expenditure of the activity. This 3D view gives a quantitative overview of how a person’s heart rate changes in response to different exercise intensity. Energy expenditure is estimated based on the horizontal and vertical displacement and the person’s weight (Weyand et al., 2010).

All the plots are generated using the plotly package, which allows for high quality and interactive visualisation, as well as ability to export and save the image.

Furthermore, CardiacProfileR can also compute several quantitative features from each heart rate profile, including resting heart rate, maximum heart rate in that period of exercise, resting indices, heart rate recovery slope, the difference between resting and ending heart rate, and coefficients from polynomials fitted to various aspects of the profile. These quantitative parameters can be extracted for downstream data analysis.

## Discussion

One novel contribution of this R package is that it turns wearable sensor data sets into high resolution quantitative phenotyping data. Data from wearable devices are beginning to be integrated into large-scale ‘omic’ data studies (Lim et al., 2018), but user-friendly bioinformatics tools that analyse these sensor data are still largely lacking. Our work therefore represents an important step towards personalised deep phenotyping (Robinson, 2012). This area of deep phenotyping research not only will fill the unmet data gap in studying the genotype-phenotype relationship, it will also provide the technological basis of potential healthcare translational software for real-time cardiovascular disease tracking systems.

## Acknowledgement

This work was supported in part by funds from the New South Wales Ministry of Health, a National Health and Medical Research Council Career Development Fellowship (1105271 to JWKH) and a National Heart Foundation Future Leader Fellowship (100848 to JWKH).

